# Disrupted activation of pain networks during attack imagery in patients with episodic migraine

**DOI:** 10.1101/2024.11.18.624121

**Authors:** Alexandre Perdigão, Inês Esteves, Ana R Fouto, Amparo Ruiz-Tagle, Gina Caetano, Raquel Gil-Gouveia, Patrícia Figueiredo

## Abstract

Migraine is a prevalent and disabling brain disorder characterized by recurrent headache attacks alternating with pain-free periods. Patients with migraine exhibit altered brain activation in response to experimental nociceptive stimuli compared to healthy controls, suggesting impaired pain inhibition mechanisms. However, the relevance of these alterations to the underlying pathophysiology of migraine attacks remains unclear. To explore this relationship, we propose a novel pain imagery paradigm aimed to induce brain activation patterns associated with migraine attacks specifically during pain-free.

The task required patients to alternate between imagining a severe migraine attack and being headache free, while healthy controls imagined severe physical head pain (e.g., toothache) and pain relief. We collected fMRI data from 14 episodic migraine without aura patients in the interictal phase and 15 healthy controls performing this task.

Both patients and controls activated pain-related brain areas during imagery of pain relative to pain relief. Moreover, controls also activated the medial pain system associated with pain inhibition and attentional modulation of pain via top-down pathways. In contrast, patients significantly deactivated these areas, namely the anterior cingulate cortex and dorsolateral prefrontal cortex. Collectively, our findings indicate altered functioning of pain networks in migraine patients, suggesting a disease-related dysregulation of pain inhibition. Eventually, the proposed attack imagery paradigm may provide a promising alternative to studies of pain mechanisms in migraine research.

## 1 Introduction

Migraine is a severe and debilitating neurological disorder characterized by recurrent attacks of severe throbbing headaches associated with symptoms such as nausea, vomiting, photophobia, phonophobia, cognitive dysfunction and mood changes (ictal phase) alternating with pain-free periods (interictal phase) [1], [2]. Current understanding suggests that migraine attacks are initiated by the activation of meningeal nociceptors and the trigeminovascular system, leading to peripheral and central sensitization [1].

Neuroimaging studies have revealed atypical activity in migraine patients compared to healthy controls in various brain regions, including those associated with pain facilitation and pain inhibition, both during [3], [4], [5], [6], [7], [8], [9], [10], [11] and between attacks [3], [4], [5], [6], [7], [8], [10], [11], [12], [13], [14], [15], [16], [17], [18], [19], [20], [21]. These studies suggest that migraine is characterized by an imbalance in pain facilitation and inhibition mechanisms, which may contribute to sensory hypersensitivity, potentially triggering attacks. Indeed, accumulating evidence indicates that migraine patients present increased sensitivity to experimental painful [3], [7], [11], [17], [18], [20], [22], [23] as well as non-painful sensory stimuli (e.g., tactile, visual, and olfactory stimuli) [24], [25], [26], [27], not only during the ictal phase [3], [7], [11], [24], [27] but also extending to the interictal phase [7], [20], [22], [23], [25], [26]. Several brain imaging studies have shown reduced habituation to experimental painful stimuli, with stronger brain activation in interictal patients relative to controls [4], [28], [29]. Nevertheless, some studies have reported the inverse relationship, mainly in regions involved in pain, inhibition such as the dorsolateral pons, anterior cingulate cortex (ACC), and dorsolateral prefrontal cortex (DLPFC) [15], [19], [21], [30].

Due to the unpredictable nature of the onset and duration of attacks, most neuroimaging studies of pain mechanisms in migraine have been conducted during the interictal phase through the application of experimental painful stimuli [31], [32]. However, somatic painful stimuli may not entirely capture the intricate pain experienced by migraine patients during an attack, which involves many other symptoms apart from pain. Therefore, it remains unknown to what extent are the previously reported changes in pain-related brain activity relevant to the pathophysiology of migraine attacks. Although a few studies have scanned patients during attacks, most recurred to pharmacological triggering to overcome the unpredictability of spontaneous attacks [9], [33], [34], [35], [36], [37], [38]. However, while nitroglycerin and nitric oxide can replicate migraine symptoms in susceptible individuals, questions remain about the extent to which these molecules contribute to the underlying pathophysiology of migraine and what influence could they have on migraine attack onset and on premonitory symptoms [32], [33], [39], [40].

Our study aimed to investigate whether pain imagery paradigms can activate the brain’s pain network in patients with episodic migraine during the interictal period. We also sought to determine whether this activation differs between patients and healthy controls. To achieve these objectives, we developed a novel pain imagery paradigm that alternates between periods of pain imagery and pain relief imagery. This approach is designed to elicit brain activation associated with migraine attacks in patients during symptom-free intervals.

## 2 Materials and Methods

This study is part of a larger research project on brain imaging in migraine (Mig_N2Treat). It employed a prospective, longitudinal, within-subject design, and consisted of a comprehensive neuroimaging protocol including arterial spin labelling MRI, diffusion MRI and several fMRI acquisitions. In this report we focus specifically on the pain-imagery fMRI data. The methods and results corresponding to the other MRI protocols are described elsewhere [41], [42], [43], [44], [45], [46]. The research protocol and statistical analysis were not preregistered. The study was approved by the Hospital da Luz Ethics Committee and all participants provided written informed consent according to the Declaration of Helsinki 7th revision.

### 2.1 Study Population

The dataset included 14 otherwise healthy adult female patients suffering from low-frequency episodic menstrual-related migraine without aura, diagnosed according to the criteria of the 3rd edition of the International Classification of Headache Disorders (ICHD-3) [47], were recruited by a neurologist during the medical appointment at the Headache Outpatient Clinic at Hospital da Luz. Besides the diagnostic criteria, other inclusion criteria consisted in: a) female; b) ages comprised between 18 to 55 years old (age 34.8 ± 8.5 years); c) at least 9 years of education and Portuguese native speaking; d) otherwise healthy (i.e., not currently diagnosed with any disease that significantly impedes an active and productive life, nor with a life expectancy below 5 years); and e) not currently receiving treatment with psychoactive drugs (including anxiolytics, antidepressants, anti-epileptics, and any drug acting as a migraine prophylactics). Exclusion criteria were: a) presence of aura; b) record of neurologic disease other than migraine; c) breastfeeding, pregnancy or after menopause; and d) any contraindications to perform MRI. Moreover, the use of contraception was not mandatory. The following clinical variables were recorded for all patients: migraine age of onset (in years); duration of migraine history (in years); migraine attack frequency (days per month); typical migraine attack duration (in hours); and typical headache intensity during migraine attacks (visual analogue scale, VAS scale). For the control group, we recruited sixteen healthy controls matched for gender, age, contraceptive use through advertisement in the general population. They met the same criteria for inclusion and exclusion, with the only difference that they did not experience regular headaches, whichever the diagnosis, and could not have any type of migraine.

The pain imagery data were collected during the patients’ interictal phase, with the absence of resolving or impending attacks confirmed 48 hours prior and 72 hours post-scan. One healthy subject was excluded due to incidental findings, resulting in a final sample of 15 healthy controls.

The final population of this study consisted of 14 female patients suffering from low-frequency episodic menstrual-related migraine without aura (age 34.8 ± 8.5 years; MIG) and 15 female healthy controls (age 30.9 ± 6.8 years; HC). The recorded demographic and clinical parameters of the participants are presented in Table 1.

**Table 1.** Demographic and clinical characteristics of the study participants: healthy controls (HC) and migraine patients (MIG). Data are expressed as mean ± SD for each group.

|  | HC (n = 15) | MIG (n = 14) |
| --- | --- | --- |
| Age (years) | 30.9 ± 6.8 | 34.8 ± 8.5 |
| Migraine age of onset (years) | | 16.8 ± 6.9 |
| Duration of migraine history (years) | | 19.7 ± 10.2 |
| Migraine attack frequency (#/month) | | 2.6 ± 1.9 |
| Typical migraine attack duration (hours) | | 40.9 ± 24.5 |
| Typical migraine headache intensity (0-10 VAS) | | 6.6 ± 1.2 |

HC = Healthy Controls; MIG = Migraine patients; VAS = Visual Analogue Scale

### 2.2 Pain-imagery Paradigm

fMRI data were acquired during the performance of a pain imagery task for 5 minutes 30 seconds, and the subjects were instructed to keep their eyes closed and received auditory cues for each task period. The pain imagery task consisted of eight cycles of alternating periods of 20-seconds of pain imagery (PI) and 20-second periods of pain relief imagery (PRI) of the corresponding pain. During the PI periods, migraine patients were asked to remain still and imagine the experience of a severe migraine attack. To achieve a comparable PI task, controls were asked to imagine a severe episode of physical head pain (e.g., tooth pain) that they had experienced. During the PRI periods, subjects were asked to imagine relief from the respective imagined pain.

### 2.3 Image Acquisition

Images were acquired on a 3T Siemens Vida MRI system (Siemens, Germany) with a 64-channel head radio-frequency coil. Structural images were acquired using a T_1_-weighted magnetization-prepared rapid gradient echo (MPRAGE) sequence (TR = 2300 ms; TE = 2.98 ms; TI = 900 ms; 1 mm isotropic resolution). fMRI data were acquired using a blood-oxygen-level-dependent (BOLD) sensitive T_2_*-weighted gradient-echo 2D-echo-planar imaging (EPI) sequence (TR = 1260 ms; TE = 30 ms; flip angle = 70°; 2.2 mm isotropic resolution; 262 volumes; in-plane acceleration with GRAPPA factor 2; simultaneous multi-slice with SMS factor 3), with each volume consisting of 60 axial slices covering the whole brain.

### 2.4 Image Analysis

#### Image Preprocessing

The imaging data were preprocessed using FMRIB Software Library (FSL) [48]. The structural images were preprocessed with nonbrain tissue removal using Brain Extraction Tool (BET) and bias field correction using FAST, and co-registered to the subject’s fMRI data space (reference volume) as well as the Montreal Neurological Institute 152 (MNI152) standard space using FLIRT and FNIRT. The fMRI data were preprocessed following a standard pipeline (as described in the literature [49], and available in https://github.com/martaxavier/fMRI-Preprocessing), consisting of the following steps: EPI distortion correction using a fieldmap image, motion correction and realignment to the middle volume (reference volume), removal of non-brain structures, estimation of motion parameters and detection of outliers, high-pass temporal filtering (0.01 Hz cut-off frequency), spatial smoothing using a 3.3 mm full width at half maximum (FWHM) Gaussian kernel, co-registration to the structural image, and subsequent normalization to the MNI152 standard space.

#### Statistical Analysis

Whole-brain voxel-wise analysis was performed on the preprocessed functional images using a general linear model (GLM) approach as implemented in FSL.

##### Subject-level Analysis

For each subject, a GLM was defined to model the PI vs. PRI BOLD response and fitted to each voxel’s time-series. The GLM included one regressor of interest, obtained by convolving the PI task timings represented by a square box function with the canonical double-gamma hemodynamic response function (HRF), along with its temporal derivative. Additionally, six motion parameters (MPs; 3 translations, and 3 rotations) and motion outliers (MOs) (identified based on a root-mean-square intensity difference between consecutive volumes above the 75th percentile plus 1.5 times the interquartile range) were included as covariates of no interest. Brain regions demonstrating PI vs. PRI activations and deactivations were identified by generating contrast maps PI > PRI and PI < PRI, respectively for each subject.

##### Group-level Analysis

All subject’s contrast maps were then entered into a group-level mixed-effects voxelwise GLM analysis. One-sample t-tests were employed to investigate brain activations and deactivations common to both groups (Brain Activations / Deactivations: HC + MIG), as well as the average activations and deactivations within each group (Brain Activations / Deactivations: HC, MIG). A two-sample t-test was used to identify regions that showed differential activation between MIG and HC groups (Differential Activation: HC vs. MIG). Whole-brain t-test images were normalized to Z scores, which were then thresholded with a voxelwise threshold of Z > 2.3, and an FWE-corrected cluster-level threshold of *p* < 0.05.

#### Relationship between Brain Activation/Deactivation and Clinical Parameters

For each cluster of brain regions exhibiting significant activation differences between patients and controls, a multivariate stepwise backward linear regression analysis was performed using the JASP software (version 0.17.2.1), with the average BOLD signal change as the dependent variable and the migraine clinical parameters (migraine age of onset; duration of migraine history; migraine attack frequency; typical migraine attack duration; and typical headache intensity during migraine attacks) as the independent variables (predictors): initially, all predictors were entered into the model and their contribution was assessed; predictors with *p* > 0.05 were progressively removed until all the predictors were statistically significant; the final model’s p-value was corrected for multiple comparisons using the Bonferroni-Holm’s method to get *p_FWE_*and considered significant if *p_FWE_* < 0.05.

## 3 Results

### 3.1 Brain Activations / Deactivations: HC + MIG

The map of brain activation / deactivation during the pain imagery task across both patients and controls (i.e., the main effect of the task regardless of the group) is presented in Figure 2. In addition, the brain regions identified in the respective clusters of voxels are listed in Table 2. When participants imagined pain (compared to imagining pain relief), certain brain regions showed increased activity (PI > PRI) in two clusters in the left hemisphere encompassing the precentral gyrus (PreCG) - involved in motor function; the inferior frontal gyrus (IFG) and supramarginal gyrus (SMG) -involved in cognitive and inhibitory control; and the postcentral gyrus (PostCG) - involved in pain sensory-discriminative processing. Moreover, decreased activity during the imagery of pain relative to relief (PI < PRI) was observed in several regions of the default mode network (DMN) – typically more active at rest, which included the posterior cingulate cortex (PCC), precuneus, angular gyrus, and medial prefrontal cortex (mPFC).

**Figure 1.**
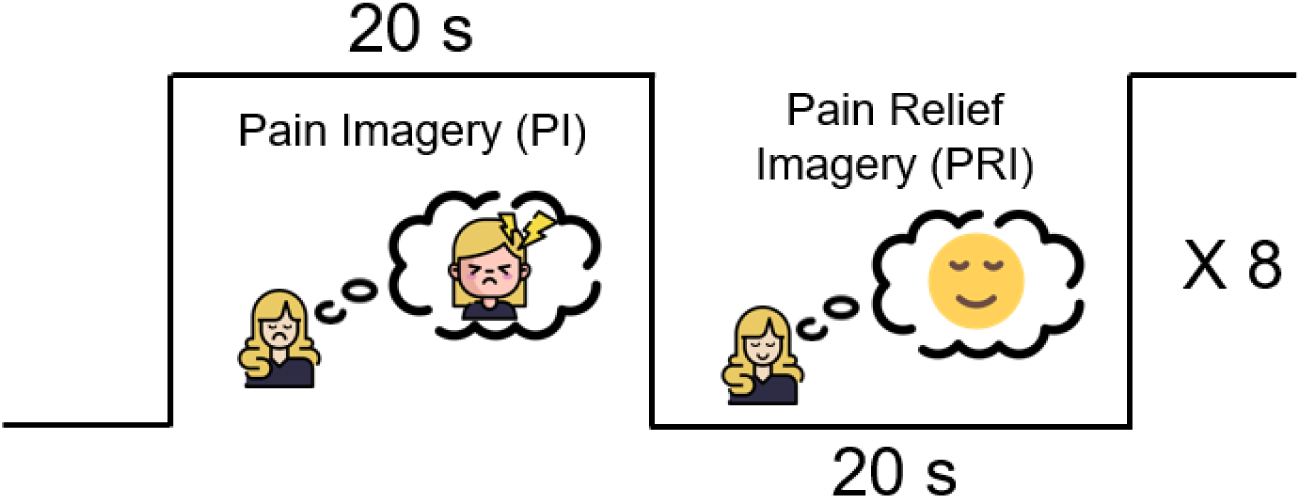
Illustration of the experimental protocol for the pain imagery task. Periods of 20s of imagery of pain (PI) are alternated with periods of 20s of pain relief imagery (PRI) of the corresponding pain, over 8 cycles, yielding a total task duration of 5 mins 30s. Subjects were instructed to keep their eyes closed, and received auditory cues for each task period.

**Figure 2.**
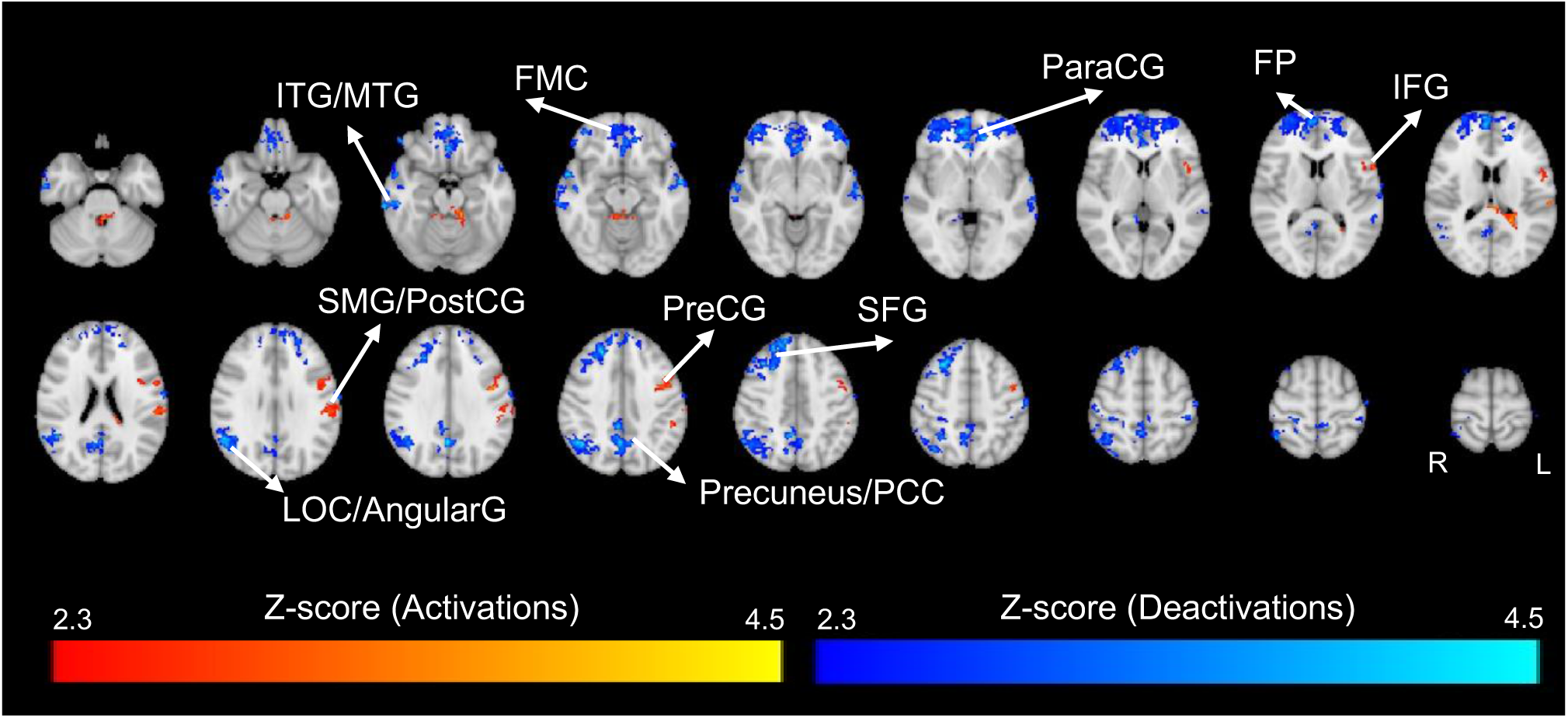
Group-level PI vs. PRI brain activations and deactivations across both groups (healthy controls and migraine patients, HC+MIG). The Z-score maps were thresholded (|Z| > 2.3; p_FWE_ < 0.05) and are overlaid on the T_1_-weighted template image in the standard MNI152 space. Acronyms: AngularG = Angular Gyrus; FMC = Frontal Medial Cortex; FP = Frontal Pole; IFG = Inferior Frontal Gyrus; ITG = Inferior Temporal Gyrus; LOC = Lateral Occipital Cortex; MTG = Middle Temporal Gyrus; ParaCG = Paracingulate Gyrus; PCC = Posterior Cingulate Cortex; PostCG = Postcentral Gyrus; PreCG = Precentral Gyrus; SFG = Superior Frontal Gyrus; SMG = Supramargial Gyrus.

**Table 2.**
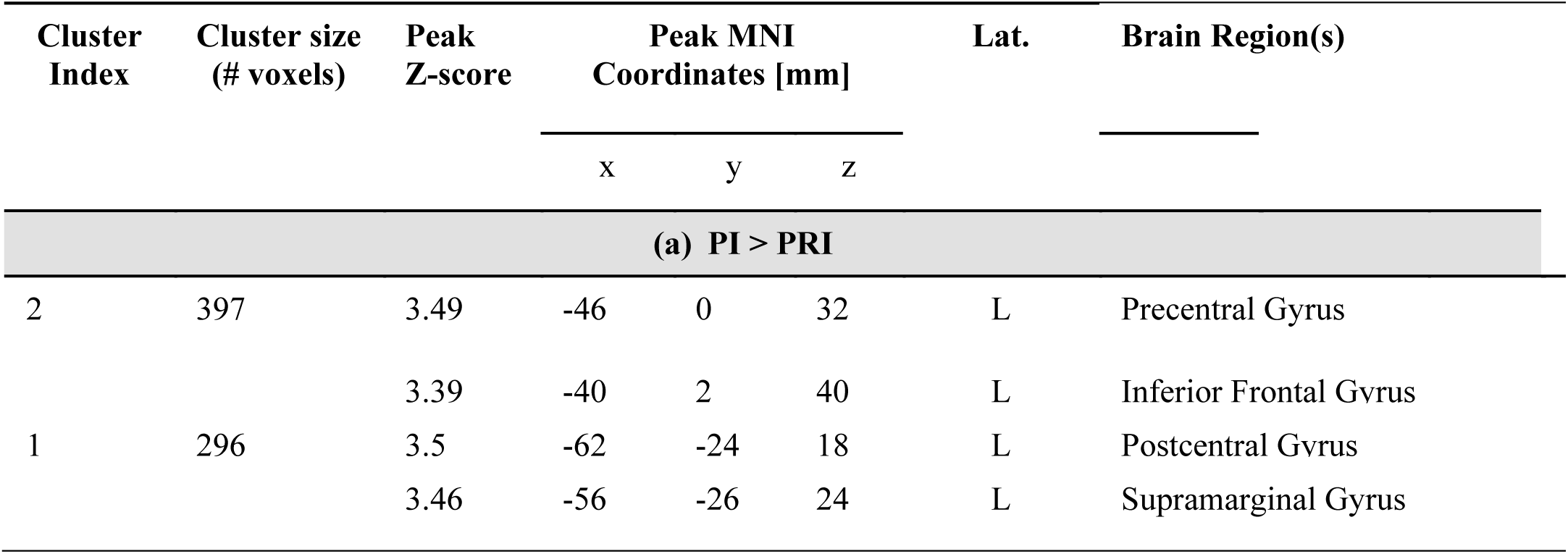
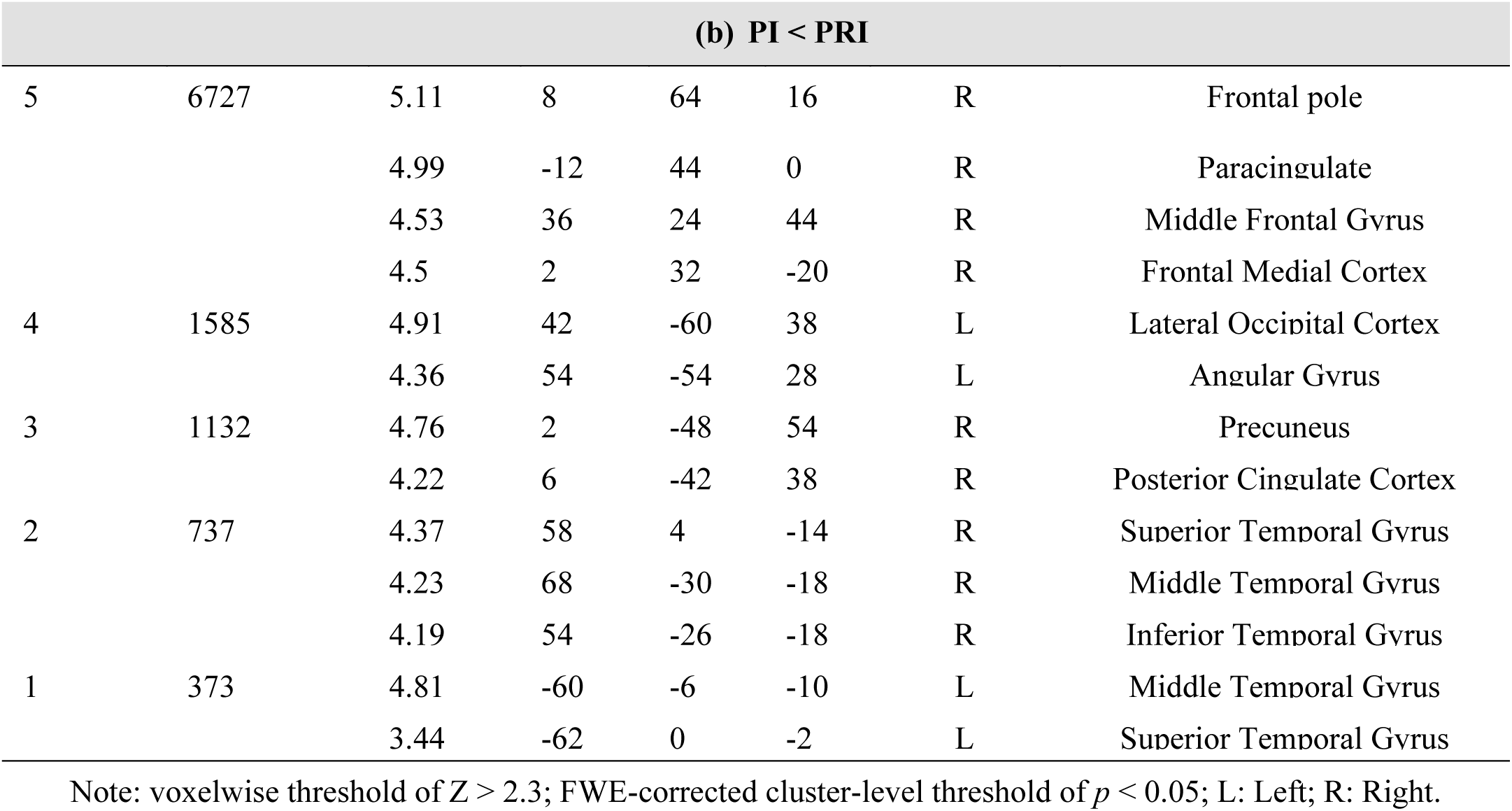
Clusters of voxels exhibiting significant brain changes across both groups: (a) PI > PRI - activations; and (b) PI < PRI - deactivations. Brain regions were obtained using the Harvard-Oxford Structural Atlases.

### 3.2 Differential Activation: HC vs. MIG

The map of differential brain activation / deactivation during the pain imagery task between patients and controls (reflecting the interaction between the task and the group) is presented in Figure 3. In addition, the brain regions identified in the respective clusters of voxels are listed in Table 3. Relative to controls, patients exhibited reduced activity (HC > MIG) in four main cluster of brain regions: (1) right DLPFC, including the frontal pole (FP) and middle frontal gyrus (MFG); (2) left DLPFC, including the IFG and the MFG; (3) ACC, paracingulate (ParaCG), and supplementary motor area (SMA); and (4) left frontal operculum cortex (FOC) and PreCG. No clusters exhibited significantly increased activity in patients compared to controls (MIG > HC).

**Figure 3.**
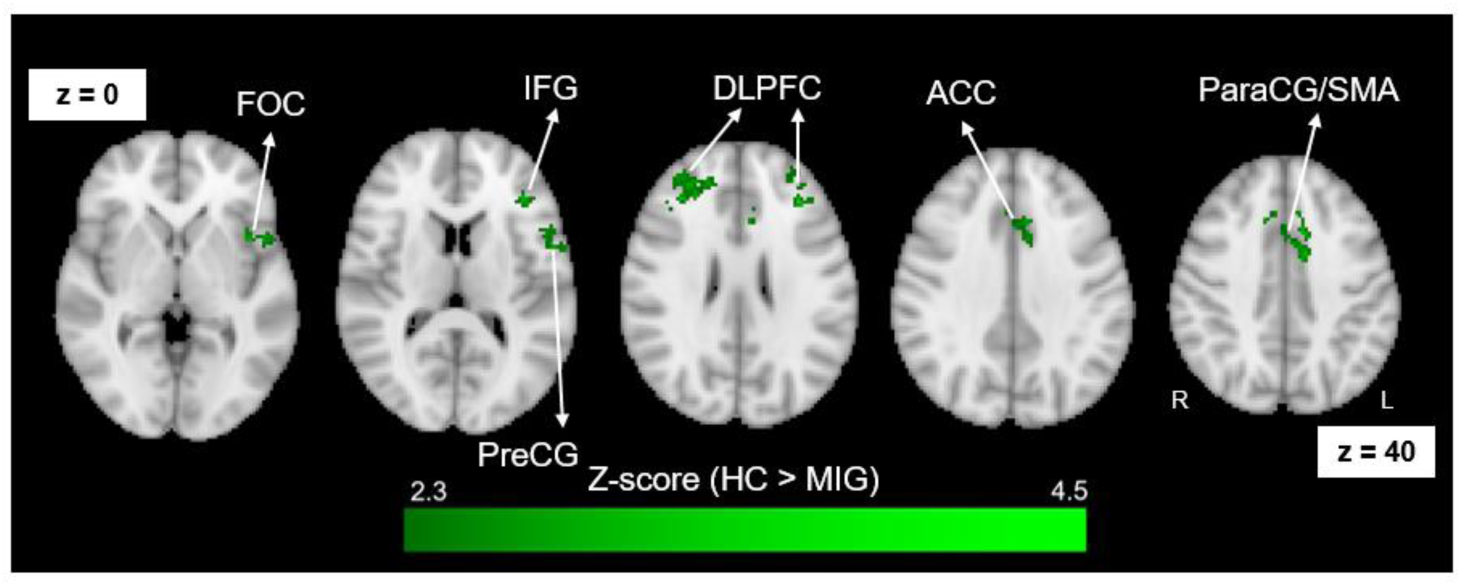
Differential brain activation during PI vs. PRI between patients and controls (HC > MIG). No significant differences were found for the HC < MIG contrast. The Z-score map was thresholded (|Z| > 2.3; pFWE < 0.05) and is overlaid on the T1-weighted template image in the standard MNI152 space. Acronyms: rDLPFC = right Dorsolateral Prefrontal Cortex; lDLPFC = left Dorsolateral Prefrontal Cortex; ACC = Anterior Cingulate Cortex; ParaCG = Paracingulate; SMA = Supplementary Motor Area; FOC = Frontal Operculum Cortex; PreCG = Precentral Gyrus.

**Table 3.**
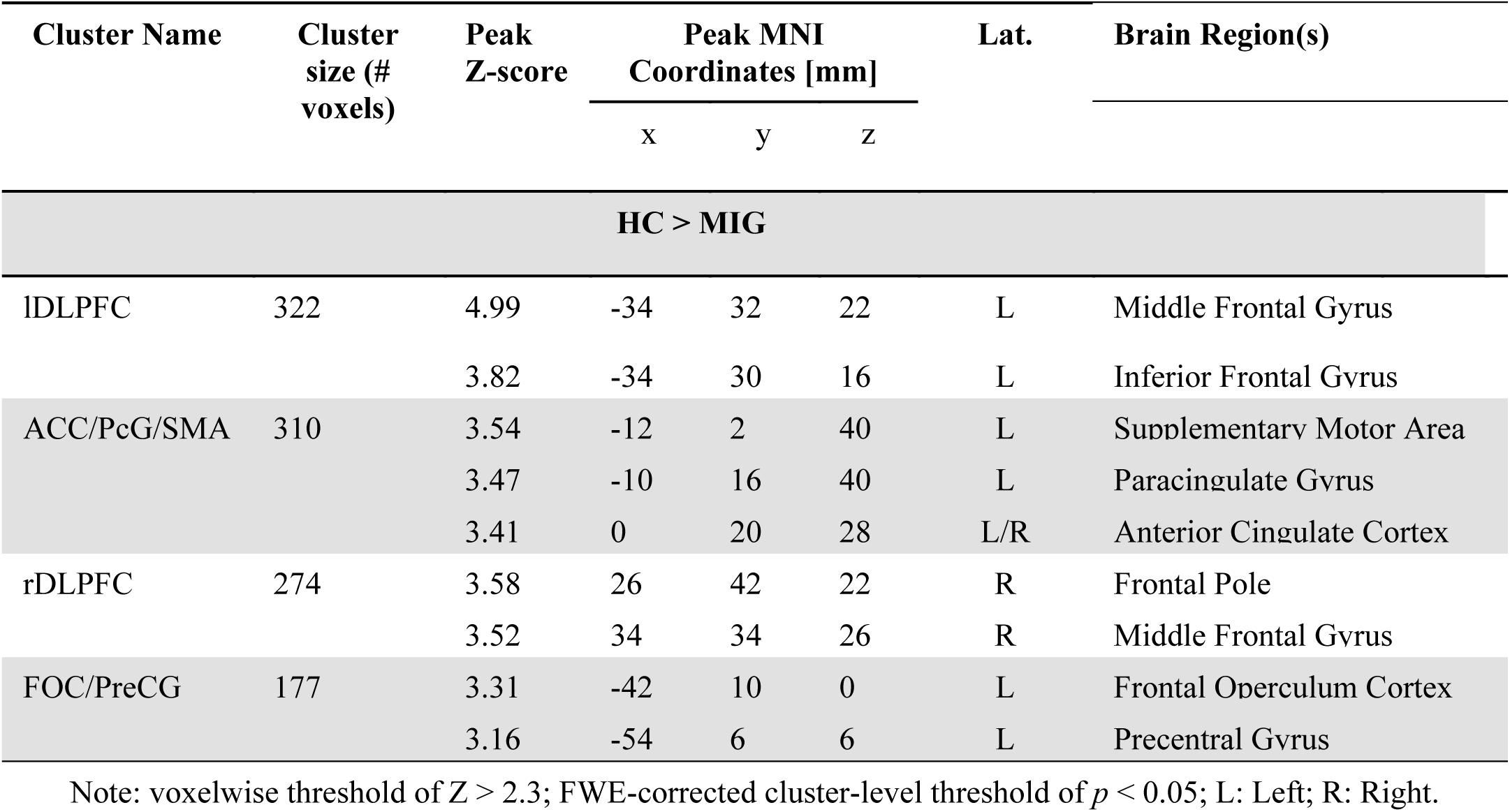
Clusters of voxels exhibiting significant brain activations and deactivations differences for the group-level HC > MIG contrast, in response to the pain imagery task. The identification of brain regions was performed based on the Harvard-Oxford Structural Atlases.

To clarify whether the observed differences corresponded to decreased activation, increased deactivation, or a change from activation to deactivation in patients relative to controls, we computed the average BOLD percent signal change (PI > PRI) in each cluster of the HC > MIG map (Figure 4). While the majority of controls activated these regions, most patients deactivated the DLPFC and the ACC, and half deactivated the IFG/PreCG. These findings indicate that these regions, normally activated by controls, became mostly deactivated in patients.

**Figure 4.**
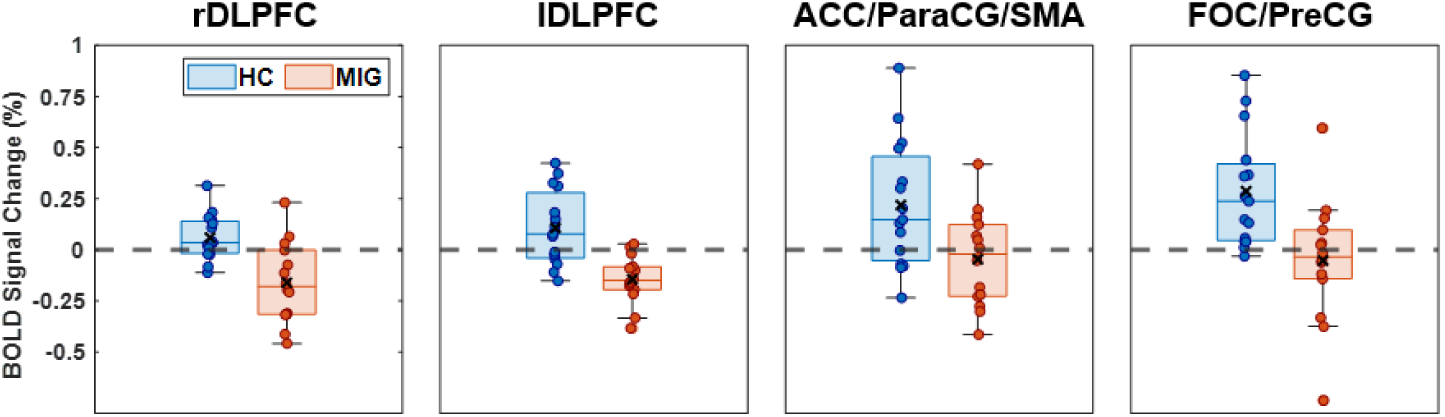
Average BOLD signal change (%) during PI vs. PRI for healthy controls (HC) and migraine patients (MIG), extracted from the four clusters of brain regions exhibiting significant differences between HC and MIG. Acronyms: rDLPFC = right Dorsolateral Prefrontal Cortex; lDLPFC = left Dorsolateral Prefrontal Cortex; ACC = Anterior Cingulate Cortex; ParaCG = Paracingulate; SMA = Supplementary Motor Area; FOC = Frontal Operculum Cortex; PreCG = Precentral Gyrus.

### 3.3 Brain Activations / Deactivations: HC, MIG

Given the significant interaction between task and group, we also present the maps of brain activation / deactivation during the pain imagery task obtained separately for patients and controls in Figure 5. In addition, the brain regions identified in the respective clusters of voxels are listed in Tables 4 and 5. When analysed separately from patients, controls also showed activations in pain-related regions, including the ACC, ParaCG, insula, PostCG, PreCG, supramarginal gyrus, IFG, superior frontal gyrus, FOC, and parietal operculum cortex. In subcortical regions, activation clusters spanned the amygdala, caudate, cerebellum, and putamen. While no significant activations were detected in the patients, they exhibited significant deactivations within the DLPFC, ACC, and ParaCG, which were activated in the controls.

**Figure 5.**
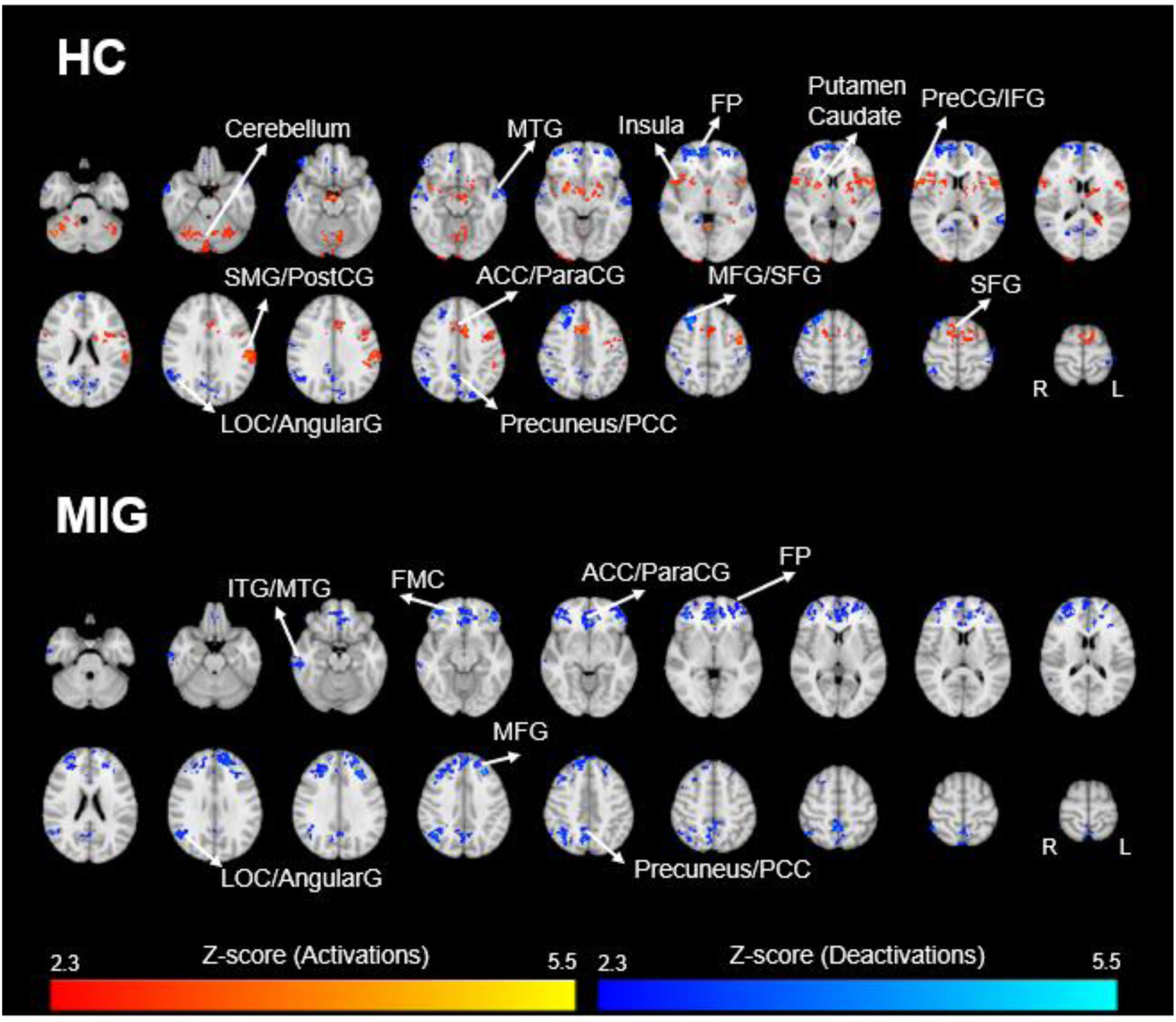
Group-level PI vs. PRI brain activations and deactivations across the HC group (top) and the MIG group (bottom). The Z-score maps were thresholded (|Z| > 2.3; p_FWE_ < 0.05) and are overlaid on the T_1_-weighted template image in the standard MNI152 space. Acronyms: ACC = Anterior Cingulate Cortex; AngularG = Angular Gyrus; FMC = Frontal Medial Cortex; FP = Frontal Pole; IFG = Inferior Frontal Gyrus; ITG = Inferior Temporal Gyrus; LOC = Lateral Occipital Cortex; MFG = Middle Frontal Gyrus; MTG = Middle Temporal Gyrus; ParaCG = Paracingulate Gyrus; PCC = Posterior Cingulate Cortex; PostCG = Postcentral Gyrus; PreCG = Precentral Gyrus; SFG = Superior Frontal Gyrus; SMG = Supramargial Gyrus.

**Table 4.** Clusters of voxels exhibiting significant brain changes in the HC group: (a) PI > PRI - activations; and (b) PI < PRI - deactivations. Brain regions were obtained using the Harvard-Oxford Structural Atlases.

| Cluster Index | Cluster size (# voxels) | Peak Z-score | Peak MNI Coordinates [mm] | Lat. | Brain Region(s) |
| --- | --- | --- | --- | --- | --- |

|  |  |  | x | y | z |  |  |
| --- | --- | --- | --- | --- | --- | --- | --- |
| (a) PI > PRI |  |  |  |  |  |  |  |
| 8 | 1161 | 4.4 | -2 | 10 | 44 | L/R | Paracingulate Gyrus |
|  |  | 4.34 | 2 | 4 | 68 | L/R | Supplementary Motor Area |
|  |  | 4.29 | 14 | 8 | 60 | R | Superior Frontal Gyrus |
|  |  | **4.25** | **-8** | **10** | **38** | **L/R** | **Anterior Cingulate Cortex** |
| 7 | 1064 | 5.38 | -52 | 8 | 12 | L | Inferior Frontal Gyrus |
|  |  | 4.56 | -46 | 6 | 38 | L | Precentral Gyrus |
|  |  | 4.55 | -44 | 2 | 50 | L | Middle Frontal Gyrus |
| 6 | 1057 | 4.86 | 46 | -62 | -28 | R | Cerebellum |
| 5 | 590 | 5.48 | 54 | 14 | 6 | R | Inferior Frontal Gyrus |
|  |  | 4.08 | 44 | 14 | -4 | R | Insula |
|  |  | 3.95 | 62 | 14 | 16 | R | Precentral Gyrus |
|  |  | 3.94 | 38 | 22 | 2 | R | Frontal Operculum Cortex |
| 4 | 499 | 5.02 | -58 | -22 | 30 | L | Postcentral Gyrus |
|  |  | 4.05 | -64 | -30 | 27 | L | Supramarginal Gyrus |
|  |  | 3.87 | -62 | -34 | 26 | L | Parietal Operculum Cortex |
| 3 | 432 | 4.85 | 22 | 4 | 6 | R | Putamen |
|  |  | 3.84 | 24 | -2 | -12 | R | Amygdala |
|  |  | 3.74 | 18 | 14 | 12 | R | Caudate |
| 2 | 421 | 3.95 | -14 | 4 | 16 | L | Caudate |
|  |  | 3.88 | -28 | -18 | 6 | R | Putamen |
| 1 | 269 | 5.28 | -26 | -66 | -22 | L | Cerebellum |
| (b) PI < PRI |  |  |  |  |  |  |  |
| 9 | 1625 | 5.14 | 30 | 58 | 0 | R | Frontal Pole |
| 8 | 889 | 5.52 | 24 | 28 | 48 | R | Superior Frontal Gyrus |
| 7 | 761 | 4.18 | 48 | -52 | 22 | R | Angular Gyrus |
|  |  | 4.15 | 46 | -66 | 40 | R | Lateral Occipital Cortex |
|  |  | 3.76 | 30 | -50 | 58 | R | Superior Parietal Lobule |
| 6 | 552 | 4.48 | -12 | -68 | 20 | L | Precuneus |
|  |  | 4.47 | -6 | -84 | 38 | L | Cuneal Cortex |
|  |  | 3.67 | -20 | -82 | 38 | L | Lateral Occipital Cortex |
| 5 | 467 | 3.96 | -60 | -20 | -4 | L | Superior Frontal Gyrus |
|  |  | 3.82 | -54 | -10 | -14 | L | Middle Frontal Gyrus |
| 4 | 442 | 5.07 | -18 | 58 | -2 | L | Frontal Pole |
| 3 | 411 | 4.59 | 70 | -22 | -2 | R | Superior Temporal Gyrus |
|  |  | 4.03 | 62 | 0 | -24 | R | Middle Temporal Gyrus |
| 2 | 274 | 4.41 | -44 | -34 | 56 | L | Postcentral Gyrus |
| 1 | 259 | 4.0 | 12 | -40 | 34 | R | Posterior Cingulate Cortex |
|  |  | 3.52 | 2 | -58 | 32 | R | Precuneus |

Note: voxelwise threshold of $Z > 2.3$ ; FWE-corrected cluster-level threshold of $p < 0.05$ ; L: Left; R: Right.

**Table 5.**
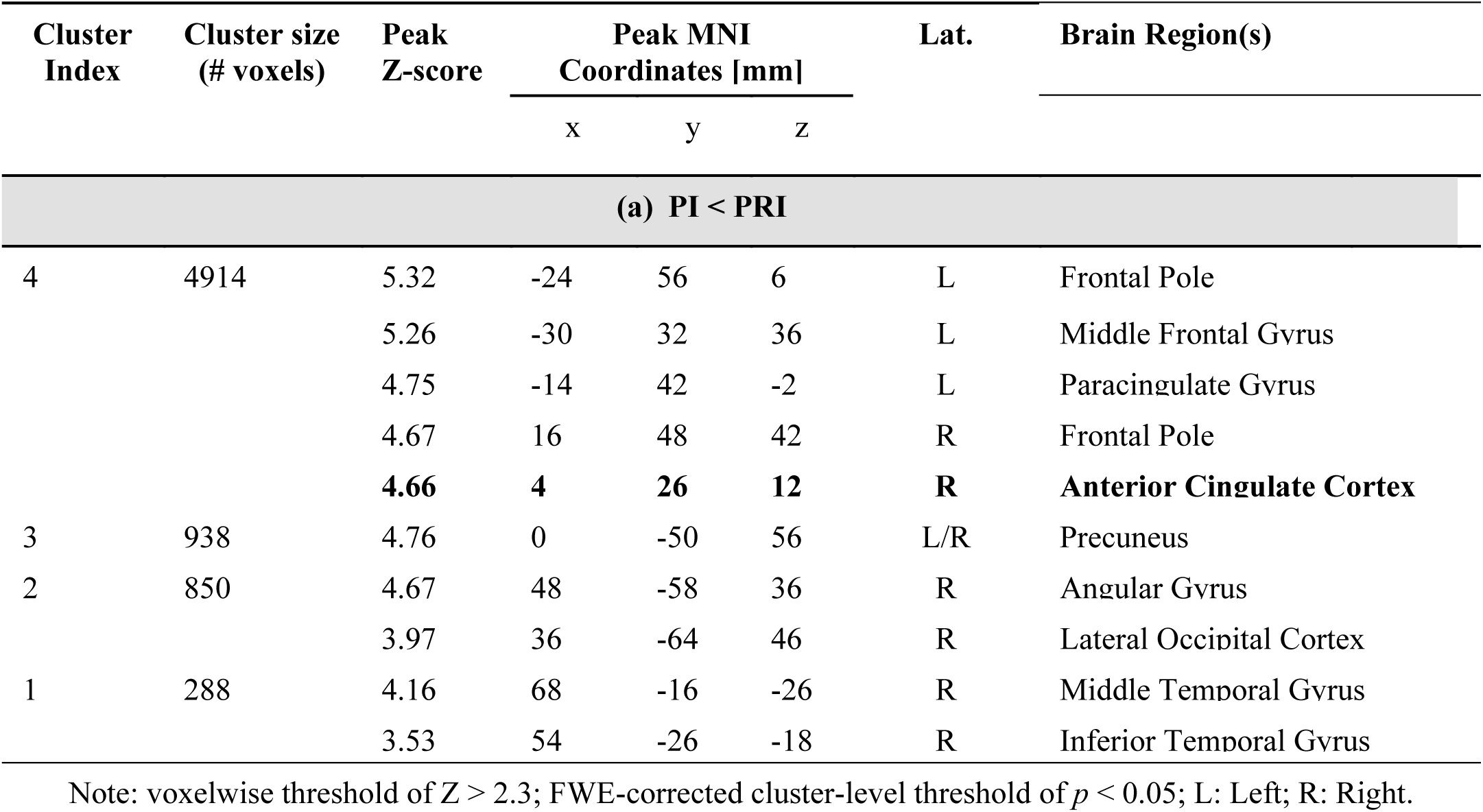
Clusters of voxels exhibiting significant brain changes in the MIG group: (a) PI > PRI – activations. No deactivations (PI < PRI) were found. Brain regions were obtained using the Harvard-Oxford Structural Atlases.

### 3.4 Relationship with Clinical Parameters

No models were found to significantly predict the BOLD signal changes extracted from the four identified functional ROIs showing the differential activations between patients and controls (HC > MIG), using migraine clinical parameters.

## 4 Discussion

In this study, we introduced a novel pain imagery paradigm that successfully elicited distinct activation patterns in pain-related brain regions in patients with episodic migraine during the interictal phase, compared to healthy controls. Our study is distinguished by its use of mental simulation of a migraine attack (pain imagery) to elicit activation of the underlying pain-related networks, without the need for invasive painful stimulation or pharmacological induced attacks, that may not fully replicate the complexity of spontaneous migraine attacks.

We found that when participants imagined pain (regardless of group) brain regions associated with movement, cognition, and pain processing showed increased activity (i.e., PreCG, IFG, and PostCG), while areas related to resting state (i.e., DMN) exhibited decreased activity. Conversely, patients demonstrated a distinct brain response to imagining pain, as they either deactivated or showed significantly reduced activation in key regions of the medial pain system - such as the ACC, DLPFC, PreCG, SMA, and FOC - compared to controls, who exhibited robust activation in these areas.

The ACC and DLPFC are key nodes of the medial pain system, involved in the cognitive, emotional, and affective aspects of pain processing, such as unpleasantness and attention to pain [50]. They play important roles in endogenous pain inhibition, attentional modulation of pain via top-down pathways, cognitive evaluation, planning motor behavior in response to nociceptive input and the emotional-affective aspects of pain perception [51], [52], [53]. Neuroimaging studies have shown that the ACC is also involved in cognitive-attentional aspects of pain and is a key cortical region that integrates the opioid-sensitive periaqueductal gray – rostroventral medulla descending modulatory system [54]. The DLPFC is also involved in cognitive-attentional and executive aspects of pain as it regulates pain-related attention, cognitive evaluation, and planning motor behavior in response to nociceptive stimuli [53]. Ultimately, the DLPFC “keeps pain out of mind”, contributing to attentional focus on pain stimuli and directing attention away from pain, mediating attenuation of pain perception through cognitive attentional control mechanisms.

The deactivation of the ACC and DLPFC in migraine patients might imply inadequate pain inhibition and dysfunctional antinociception. Our findings are consistent with prior research using painful stimulation. One study applied repetitive trigeminal stimulation and reported decreased activation of the DLPFC, ACC, red nucleus, and ventral medulla in migraine patients, whereas controls exhibited increased activation in these regions with such stimulation [21]. Another study found a significant reduction of the BOLD signal in the pons in response to painful stimulation among during the interictal phase. This region contains the nucleus cuneiformis, a predominantly inhibitory region in the pain descending pathways, supporting decreased pain response inhibition in migraine [19].

Previous studies reporting decreased activity of the ACC in migraine patients indicate a possible relationship between deactivation of the ACC and migraine attack frequency, suggesting that recurrent attacks lead to progressively more abnormal ACC activity. On the other hand, it could also reflect an abnormal baseline activity predisposing patients to more frequent attacks. Russo and colleagues [15] found a significant negative correlation between ACC BOLD signal changes in response to trigeminal stimulation and migraine attack frequency, i.e., the higher the attack frequency the greater the ACC deactivation. Another study, using magnetoencephalography identified a negative correlation between attack frequency and node strength in the bilateral ACC among migraine patients [55]. In a structural imaging study, Valfrè et al. (2008) [56] showed that migraine patients, compared to controls, exhibited reduced ACC grey matter volume, which positively correlated with the frequency of migraine attacks.

A magnetoencephalography study found reduced beta oscillatory connectivity in the ACC was linked to migraine chronification, highlighting the potential role of ACC dysfunction in increasing attack frequency [55].

In our study, deactivations were also found in the PreCG, SMA, and FOC in migraine patients compared with controls. These regions play distinct yet interrelated roles in motor control and cognitive processes such as motor planning, coordination, initiation, and response inhibition of motor actions, as well as in sensory-processing of pain [57]. This result is consistent with previous findings by Schwedt et al. (2014) [18], who reported less pain-induced activation in migraineurs compared to controls in similar regions, including the PreCG and SMA. The authors suggest that activation in these regions may indicate a preparation for moving the body away from painful stimuli. Since physical movement worsens the pain during migraine attacks, decreased activation in these regions might suggest that migraine patients learned to avoid movement during the pain associated with attacks.

To our knowledge, this is the first study using a pain imagery paradigm to study migraine associated brain activity using fMRI. Previous studies using pain imagery tasks have elicited pain imagery using pictures [58], [59] or imagination of pain [60], [61], [62] and were able to activate pain related brain regions. These studies show that pain imagery activates a network of brain regions that largely overlaps with the network activated during physically induced pain and suggest a possible distinction in how pain imagery engages the sensory and affective components of pain processing. Supporting out findings, while imagined pain consistently activates areas associated with the affective dimension of pain (e.g., ACC, PFC, and anterior insula) [58], [59], [60], [61], [62], activation in sensory discriminative regions (e.g., primary and secondary somatosensory cortices) is more variable [58], [62].

If migraine attack imagery proves able to consistently elicit activation of the underlying pain neural networks, it may provide a powerful, non-invasive, ecologically-valid tool for research of migraine pain episodes that is more specific than experimental painful stimulation and possibly is closer to the individual experience of the ictal phase, overcoming some of the logistical caveats associated with studying the unpredictable ictal phase or resorting to pharmacological attack induction with inherent risks and patients discomfort.

### Limitations

This study presents limitations. Studies with larger sample sizes are needed to replicate our findings and enhance the robustness of group-level inferences and correlation analyses. Despite the relative simplicity of studying patients in the interictal phase, longitudinal studies that track patients across the entire migraine cycle could provide further insights into the evolution of brain alterations leading up to a migraine attack. It would also be important to include high-frequency and chronic migraine patients in future studies, to investigate changes associated with the progression of the disease. While promising, the pain imagery paradigm requires further validation to ensure its effectiveness in capturing pain-related networks. Improved experimental designs could be developed to better simulate the complexity of a spontaneous migraine attack, for example by combining pain imagery with trigeminal nociceptive stimulation. Furthermore, questionnaires assessing the vividness of pain imagery could be implemented to determine the participants’ ability to perform the task effectively.

## 5 Conclusions

This works demonstrates that a pain imagery paradigm is capable of eliciting differential activation in relevant pain-related brain regions among individuals with episodic menstrual migraine during the interictal phase. This finding holds great promise for future applications in migraine research, as it offers an alternative to painful stimulation or pharmacological attack triggering that is potentially closer to the experience of migraine pain, while overcoming the challenges of studying spontaneous migraine attacks. By employing the proposed pain imagery paradigm, we obtained valuable insights into migraine pathophysiology, by providing imaging evidence of altered functioning of brain regions associated with pain processing in the migraine brain.

## Acknowledgements

This work was supported by LARSyS FCT funding (DOI: 10.54499/LA/P/0083/2020, 10.54499/UIDP/50009/2020, and 10.54499/UIDB/50009/2020), and FCT grants PD/BD/150356/2019, PTDC/EMD-EMD/29675/2017, LISBOA-01-0145-FEDER-029675.

## Conflicts of interest

The authors have no conflicts of interest to declare.

## Notes

### Competing Interest Statement

The authors have declared no competing interest.

